# Ontogenic shift in the usage and function of a symbolic bird gesture

**DOI:** 10.64898/2026.01.07.698152

**Authors:** Mylène Dutour, Valentine Ouvry, Thierry Lengagne, Cédric Girard-Buttoz

## Abstract

Gestures are an integral part of human communication, and the characterisation of pointing (deictic) and meaningful (symbolic) gestures in non-human primates helps retracing the origins of our own communication system. A recent discovery in Japanese tits (*Parus minor*) suggested that individuals use a symbolic wing-fluttering gesture to prompt the mating partner to enter the nest first. Symbolic gestures are thus likely more ancestral than previously thought, but the presence of this gesture in related species, the degree to which it is tied to feeding context and whether gestural usage changes across development remain unexplored. To fill these gaps, we studied the wing-fluttering gesture in great tits (*Parus major*), observing both fledglings and adults in various contexts around nests and away from nests in their natural environment. Our findings showed that wing-fluttering was primarily used by females near nests, almost exclusively in the presence of their mate, and stopped when the mate entered the nest. As in Japanese tits, our results suggest that wing fluttering serves as a symbolic gesture prompting the mate to enter the nest more quickly to feed the nestlings. In contrast, fledglings used wing-fluttering to prompt their parents to feed them and stopped gesturing after having been fed. Our study further indicates deep evolutionary roots of symbolic communication and highlights how functional shifts in gestural usage can operate across development to match age-specific needs.

## Introduction

Gestural communication plays a central role in the social lives of many animal species and offers key insights into the evolutionary roots of complex communication systems, including human language (Pika et al., 2005; Moore, 2016; Eleuteri et al., 2025). To explore these roots, comparative researchers mostly focus on our closest living relatives, the non-human primates. Pre-linguistic children are used as a point of reference in such studies, as they use a variety of gestures from an early age, such as showing or requesting objects, waving goodbye, or pointing to elements in their environment (Bates et al., 1979; Liszkowski et al., 2004). As a result, the criteria used to identify gestures in pre-linguistic children have been widely applied in studies of gestural communication in non-human primates (Leavens et al., 2005; Moore, 2016; Pika et al., 2005).

Currently, there is considerable variability in how gestures are defined across studies and species. Here, we adopt a definition in line with established criteria: to qualify as a gesture, individuals must exhibit a specific body movement only in the presence of the target receiver, this movement should elicit a specific behavioural response in the receiver and stop once the receiver has produced this particular behavioural response (Hobaiter and Byrne, 2014; Pika and Bugnyar, 2011; Vail et al., 2013). Since we focus on the visual modality - excluding tactile gestures such as ’touch’ or ’push’ - a final criterion is that the movement must not act as a direct physical agent on the receiver.

Gestures can be classified into two categories: deictic gestures (such as pointing to objects) and symbolic gestures (which convey specific messages) (Goodwyn et al., 2000). Evidence is steadily accumulating for deictic gestures in a wide range of taxa (Pika and Bugnyar, 2011; Vail et al., 2013; Eleuteri et al., 2025), suggesting a deep evolutionary origin of this type of gestures. In contrast, reports of symbolic gestures remain limited to a much narrower range of species. Symbolic gesturing has so far been found almost exclusively in some primate species (great apes; Kalan et al., 2025), including humans (Goodwyn et al., 2000). Outside of primates, such gestures have been anecdotally reported in the orange-winged Amazon (*Amazona amazonica*) (Moura et al., 2014), and was recently shown in a single bird species; the Japanese tit (*Parus minor*) (Suzuki and Sugita, 2024). In this species, females use wing-fluttering to prompt their mates to enter the nest cavity first to feed the nestlings. Wing-fluttering appears to function as a symbolic gesture conveying an ‘after you’ message (Suzuki and Sugita, 2024). However, this study was based on a relatively small number of observations, and recent analyses have revealed high variability in responses, raising questions about the generalizability of the findings (Da Silva and Matsushita, 2025). To tackle these limitations, an evaluation of the presence and usage of this gesture with a larger sample size may allow us to investigate the inter-individual variability. Another limitation of the study is that it relied solely on observations of birds at the nestbox, impairing the ability to assess whether the wing-fluttering gesture is used in different situations and has flexible functions across contexts.

Another important aspect omitted in the study on Japanese tits is the ontogeny of this communicative signal. This assessment is important because, in humans, language is a learned communicative system in which the form of arbitrary signals needs to be matched to a specific function, often unrelated to the actual form of the signal. Such a *form to function mapping* is acquired throughout development, and functional usage of signals can change across development. Gestures in particular exhibit a flexible relationship between signal forms and communicative functions, where different gestures can be used for the same goal, and the same gesture can be used for different goals (e.g., Pollick and de Waal, 2007).

Interestingly, we observed wing-fluttering behaviour in fledglings of several bird species (own unpublished data), where fledglings performed the gesture in front of their parents prior to being fed, in a manner similar to how adult Japanese tits use wing-fluttering when they prompt their mates to enter the nest cavity first to feed the nestlings. Although the immediate function appears different - soliciting food for themselves versus prompting a mate to feed nestlings - the gesture occurs in a similar context of food provisioning. This raises the possibility that the same gesture can serves different communicative functions over the course of development, suggesting a degree of flexibility in gesture use.

In this study, we investigate whether the closest phylogenetic relatives of the Japanese tit, the great tit (*Parus major*), use gestures that could qualify as symbolic, aiming to clarify their functional consistency and contribute to a broader understanding of gestural communication in birds, helping to trace the evolutionary roots of our own language. We also examine the functional use of this gesture in fledglings to investigate the potential developmental shift in its function between fledglings and adults. Great tits are socially monogamous birds that nest in tree cavities, similar to Japanese tits. Both species present a highly similar vocal communication system with the same single call types and the same compositional combination of two call types into ordered sequences (the “ABC-D” alert-recruitment call sequences) (Suzuki et al., 2016; Dutour et al., 2019a; Suzuki and Matsumoto, 2022). The study of their vocal communication, in particular the detailed understanding of the ABC-D sequence (Schlenker et al., 2024), provided a valuable window into the evolutionary origins of structural complexity in human language. Here we expand on these findings by characterising similarities and differences in the use of a gesture that qualifies as symbolic. Symbolic gestural and verbal signals are abundantly used in our daily communication and are fundamental to language, and our study helps trace the evolutionary origin of such signals and of their ontogeny.

In our study, we collected data on wing-fluttering contextual usage and behavioural outcomes at the nestboxes of 15 pairs of great tits, as well as on 83 adults and 24 fledglings in their natural environment (natural nest holes or away from the nest). Using these detailed observations, we evaluated whether wing-fluttering also met the criteria of a symbolic gesture and whether functional usage shifted across development.

## Methods

### Study site and observations of adults at the nestboxes

We conducted this study in a mixed deciduous-coniferous forest near Lyon, France (4.91°E, 45.95°N), where 120 nestboxes, installed in 2016 at ∼1.5 m height on tree trunks and spaced at least 50 m apart (Dutour et al., 2019b), have been maintained since. Nestboxes were visited at least every two days from the beginning of the breeding season 2025 (late March) to determine laying and hatching dates. For behavioural data collection, we selected nestboxes occupied by great tits that met the following criteria: (i) nestlings were 8-16 days old during the observation period to be in line with the previous study of Suzuki and Sugita (2024); (ii) the nestbox had to be sufficiently unobstructed to allow clear observation from a distance of ∼20 m by the three observers; and (iii) the surrounding area within a 5-m radius included small bushes or shrubs where great tits could perch before entering the nestbox. Among the 19 selected nestboxes, four were excluded from the analyses: one because no observations could be made, as the parents remained beyond the 5-m radius from the nestbox (see below for details); one because only a single individual was observed within the 5-m radius and its sex could not be determined due to an atypical breast stripe pattern combined with very shy behaviour; and two because the nestboxes were predated a few hours or days before the observations. For three additional nestboxes, although the parents approached within the 5-m radius, no feeding of nestlings was observed during the observation period.

Observations were conducted from April 25 to May 14, 2025 between 9am and 12pm (Central European Time) under calm and dry weather. Three observers, concealed within the vegetation and positioned at a distance of approximately 20 m from each nestbox (mean ± SD = 20.7 ± 3.3 m; range: 15-31 m), observed the breeding pair for an average of 145 min (range: 135-155 min). The 20 m distance was chosen to minimize disturbance to the tits and in line with a previous study on Japanese tits in which a 15-m distance was used (Suzuki and Sugita, 2024). Prior to each observation session, the three observers spent 15 min in the concealed position (pre-observation period) to allow the birds to habituate to their presence.

Each time one of the parents approached within a 5-m radius of the nest constituted a separate observation. During each observation, we noted the social context, distinguishing two social contexts: (i) the bird arrived alone and did not encounter its partner before entering the nest, or (ii) the bird encountered its partner near the nest before entering. For each observation, we also recorded the sex of the individuals, whether they performed wing-fluttering behaviour, the time elapsed between their arrival within the 5-meter radius and their entry into the nest, and if they entered in the nest or not.

A total of 893 observations were recorded across 15 nestboxes (N = 30 individuals; 15 males and 15 females). We excluded 110 observations; 93 for which we were unable to determine the sex of the individual, 14 for which we could not determine whether the gesture was performed, and 3 for which we could not determine either of the two parameters. As a result, we retained a total of 492 observations with nest visitations and 291 observations without nest visitations (i.e., cases where individuals entered the observation zone but left without feeding) for analysis. We obtained 121 observations (112 for males and 9 for females) in which both individuals were present and for which we were able to calculate the latency to enter the nestbox.

During observations, each observer used binoculars and a voice recorder to verbally describe in real time everything they observed, allowing for detailed and accurate behavioural annotation. Two observers, positioned together on one side of the nestbox, focused on one parent each, while a third observer, positioned on the opposite side, monitored the overall activity around the nestbox to help capture behaviours that might otherwise be missed and to provide contextual information. The final dataset consisted of the most reliable observations confirmed by the three observers and the recordings; the agreement among the three observers was very high (∼ 99%) for all variables. We also filmed a few minutes of observations for 8 nestboxes.

To confirm the validity of our behavioural classification (i.e., wing-fluttering vs. no wing-fluttering), we conducted a test with nine adult people who were not ornithologists and were unaware of the study’s hypothesis. First, two ∼10-sec video clips were shown as examples, each illustrating a behaviour we had previously classified as either wing-fluttering or not. Then, participants were shown six new randomly selected ∼10-sec clips (three showing wing-fluttering and three without) and asked to classify whether bird exhibited wing-fluttering in each. The classifications made by these hypothesis-blind observers showed strong agreement with our own assessments, with an accuracy rate of 92.6% (50 correct classifications out of 54 total responses), ensuring that there was no bias in our behavioural recording.

### Observations of adults and fledglings away from nestboxes

To determine whether 1) adult great tits exhibit wing-fluttering in other contexts than nestling provisioning at nestboxes, and 2) fledglings exhibit wing-fluttering in front of their parents before being fed, in a manner similar to how adult tits use wing-fluttering when they prompt their mates to enter the nest cavity first to feed the nestlings, we conducted additional focal observations from 5 to 26 June 2025. These observations were conducted independently of the nest-based recordings and were carried out away (∼ 20 km) from the monitored nestboxes to limit nest-related behaviours. Moreover, to minimize disturbance and reduce the potential impact of human presence on the behaviour of the focal individual (given that, unlike nestbox observations where observers were hidden in vegetation several ∼20 meters away, these observations required closely following great tits; within ∼10 meters), all these focal observations were conducted by a single observer. To locate great tits, the observer walked slowly along trails in large urban parks or mixed forests, in areas where we had previously studied tits a few years earlier (Dutour et al., 2017; Dutour et al., 2019a). Juveniles are distinct from adults and easily recognisable due to being duller in coloration, with a less pronounced black belly stripe, and pale-yellow cheeks.

Individuals were selected opportunistically and followed with binoculars for focal observations until they could no longer be observed. Each observation involved continuous monitoring of a single individual in its natural environment. The observers recorded spontaneous the bird’s behaviours in the field as they naturally occur. During each observation, we recorded whether the individual exhibited wing-fluttering behaviour, the context, and the number of conspecifics or heterospecifics present within a 5-meter radius. Whole observation sessions were audio-recorded using a dictaphone, with the observer verbally describing all visible behaviours in real time. In total, we observed 107 individuals (42 females, 41 males and 24 fledglings), with a cumulative observation time of 11 hours (an average of 6 min of continuous focal observation per individual; range: 1-25 min) and 225 observations over 13 days. Out of 83 observed adults, 5 individuals (2 females and 3 males) were observed near their natural nesting cavities (i.e., in natural settings, not nestboxes). Since individuals in the population are unringed, we maintained a minimum distance of 200 meters between individuals of the same sex to minimize the risk of observing the same individual more than once (Salis et al., 2022).

### Statistical analysis

We ran a series of Bayesian Generalised Linear Mixed Models (GLMM) to test our predictions on the function of the wing-flittering gesture in great tits. The first series of Models analysed the bird behaviours at the nestbox, and we tested the effect of sex and mate presence on the production of wing-fluttering (Model 1) and the effect of wing-fluttering on which sex entered the nestbox first (Model 2), as well as on how fast the partner entered the nest (Model 3). To confirm that wing fluttering can be defined as a gesture in these birds, we tested one key component of gestures, namely that the gesture ends when the goal is reached. We assessed whether the birds were more likely to stop than to continue gesturing after the partner entered the nestbox in a fourth Model (Model 4). The gesture was observed only once across all observations of adults away from the nest cavity (40 females and 38 males observed). Therefore, no statistical analyses were performed, and only descriptive data are presented.

We also ran two additional analyses to evaluate the function of the wing-fluttering gesture in fledglings observed in the wild away from the nestboxes. Since we hypothesised that wing fluttering in fledglings is directed towards their parents, we tested the effect of parent presence on the likelihood of producing the gesture (Model 5). We also tested whether fledglings were more likely to stop rather than continue gesturing once they had been fed (Model 6). We ran our models using the function brm from the package ‘brms’ (Buerkner, 2018) in R (version 4.4.2), which provides an interface with STAN (Stan Development Team, 2020).

#### Model 1: Social influence on the wing-fluttering pattern of great tits

In our first model (Model 1), we used a GLMM with a “bernoulli” error distribution and a logit-link function to model the effects of individual sex and mate presence on the production of gestures around the nestboxes. Each observation for each individual constituted a datapoint, and we used whether the individual gestured (Yes/No) as the response variable. We used the individual sex and mate presence (present or absent) as two categorical predictors. We also modelled the interaction between sex and mate presence to account for the possibility that the social function of the gesture is sex dependent. Since we recorded several observations on the same individuals and since two individuals were observed per nest, we also included individual ID and nest ID as random effects in the model.

#### Model 2: effect of wing-fluttering production on the sex of the first to enter the nestbox

In our dataset, males gestured at the nest only 3 times out of 170 observations when both mates were present, which did not allow us to test the effect of male gesturing on the sex of the individual that first fed the youth. Accordingly, in our second model (Model 2), we analysed the effect of female gesture on this likelihood and discarded the observations in which males gestured. In this model, we used only observations in which both males and females were present at the nest simultaneously. Each observation constituted a datapoint. We used a “bernoulli” error distribution and a logit link function to model the effect of female gesture production on whether the male or the female entered the nestbox first (binomial response, did the male enter the nestbox first (Y/N). We used female gesture (Y/N) as a categorical predictor and controlled for the sex of the individual who arrived first at the nest since this could influence who entered the nestbox first. In Model 2, we also accounted for multiple observations at the same nest (i.e., of the same pair of individuals) by including nest ID as a random effect.

#### Model 3: effect of female gesture on the latency for males to enter the nestbox

For the same reasons as for model 2, related to limited sample size (N = 9 observations in which females entered the nestbox, the male was present, and we could record the latency to enter the box), we focused on female wing-fluttering when assessing the impact of this body movement on the latency for the mate to feed the nestlings. We used another GLMM (Model 3) with a Weibull error distribution and log-link function. Each observation where both mates were present at the nestbox and for which male entered the nestbox, and we had information on whether the female gestured or not, constituted a datapoint. We used female gesture (Y/N) as a test predictor and controlled for whether the male or the female arrived first at the nest. To account for repeated observations of the same pair of individuals at the same nest, we included nest ID as a random effect in Model 3.

#### Model 4: Influence of male behaviour on gesture interruption in females at the nestbox

In a fourth model, we aimed to verify one key element of what constitutes a gesture, namely that when the goal communicated by the gesture is achieved, the individual stops gesturing. Since the after-you gesture has been shown in another tit species to promote the feeding of the young by the other individual of the couple, we hypothesised that the function in the great tit would be the same. We used an intercept-only GLMM with a “bernoulli” error distribution and a logit link function to model the likelihood of female tits to stop gesturing when the male started to feed the young. Each wing-fluttering produced by a female constituted a datapoint, and the response variable was whether the female stopped wing-fluttering when the male started to feed the young (Yes or No). This model did not comprise any predictors, but we included female ID as a random factor to account for repeated observations of gestures from the same female.

#### Model 5: Social influence on fledgling gesture production

After testing the social triggers and consequences of wing-fluttering produced by adults, we focused on the production of the same gesture by fledglings. We first tested whether wing fluttering is more often produced when the hypothesised targets of the gesture, the parents, are present. We used a GLMM with a “bernoulli” error distribution and a logit-link function to model the effect of parent presence (binomial predictor with presence Y/N) on the production of gestures by fledglings (Model 5). Each observation for each individual constituted a datapoint, and we used whether the individual gestured (Yes/No) as the response variable. We recorded several observations on the same individuals within the same observation session, as we split the data into separate observations every time the parents left and returned. To account for these repeated observations, we included fledgling ID as a random effect in the model.

#### Model 6: Influence of parent’s feeding behaviour on gesture interruption in fledglings

Since fledglings do not feed other conspecifics, we hypothesised that the wing fluttering gesture produced by fledglings was directed at parents and aimed at receiving food from them. Therefore, similar to Model 4, we used an intercept-only GLMM with a “bernoulli” error distribution and a logit link function to model the likelihood of fledglings to stop gesturing when one of the parents fed them Each gesture produced by fledglings constituted a datapoint, and the response variable was whether the fledglings stopped gesturing when the parent started to feed them (Yes or No). This model did not comprise any predictors, but we included fledgling ID as a random factor to account for repeated observations of gestures from the same fledgling.

For all models, we included the maximum random slope structure of the fixed predictor within each random effect. We ran 4000 iterations (2000 for warmup) on 4 chains. We used weakly regularising priors for the fixed effects (Normal (0,1)) and the priors given by default by the function “get_prior” of the package “brms” for the random effects (i.e., Student t (3, 0, 2.5) for the random intercepts and slopes). We chose weakly regularising priors for the fixed effects since they give less weight to outlier data points and help constrain model predictions to biologically meaningful estimates and credible intervals (Lemoine 2019). For Model 5, however, those priors resulted in a poor model fit with a low effective sample size, even when multiplying by 4 the number of chains and iterations. This could be related to the fact that gestures were only observed in the presence of parents, and to the effect of parent presence that was very large. Accordingly, for this model, we used the default flat prior of brms, which provided a much better fit. We then extracted the 95% and 89% CIs for each fixed effect from the model’s posterior distribution.

Sampling diagnostics (Rhat = 1 for all predictors in all models) and trace plots confirmed chain convergence for all models. Effective sample sizes (all > 4000) confirmed no issues with autocorrelation of sampling for all models. Note that the effective sample size is a measure of autocorrelation and does not correspond to the number of data points that were used for the model. We provide the posterior predictive check of each model in Figures S1-S6. In the result section, we provide the proportion of the posterior distribution in support of the direction of a given coefficient as *P.support*.

## Results

### Social triggers of gesture in adults at the nestboxes

In Model 1, (N = 783 datapoints from 567 independent observations on 29 birds (14 females and 15 males) from 15 nests), we found strong statistical support for an effect of both individual sex (ß= -1.78, 95% CI [-3.04, -0.57], P.support = 0.997) and mate presence (ß= 1.91, 95% CI [ 0.65, 3.09], P.support = 0.998) on the likelihood for the bird to produce a gesture at the nestbox (Table 1, Fig. 1). In contrast, we did not find statistical support for the interaction between sex and mate presence to impact the likelihood of gesture (89% CI overlapped 0 and P.support = 0.844, Table 1). Birds from both sexes were 15 times more likely to gesture in the presence than in the absence of mates (gestures were observed in 93 of 432 observations = 21.52% in the presence of a mate vs. in 5 of 351 observations = 1.42% in the absence of a mate, Fig. 1). Stated otherwise, 94.5% of gestures (93 out of 98) were observed when the mate was present.

**Fig. 1.**
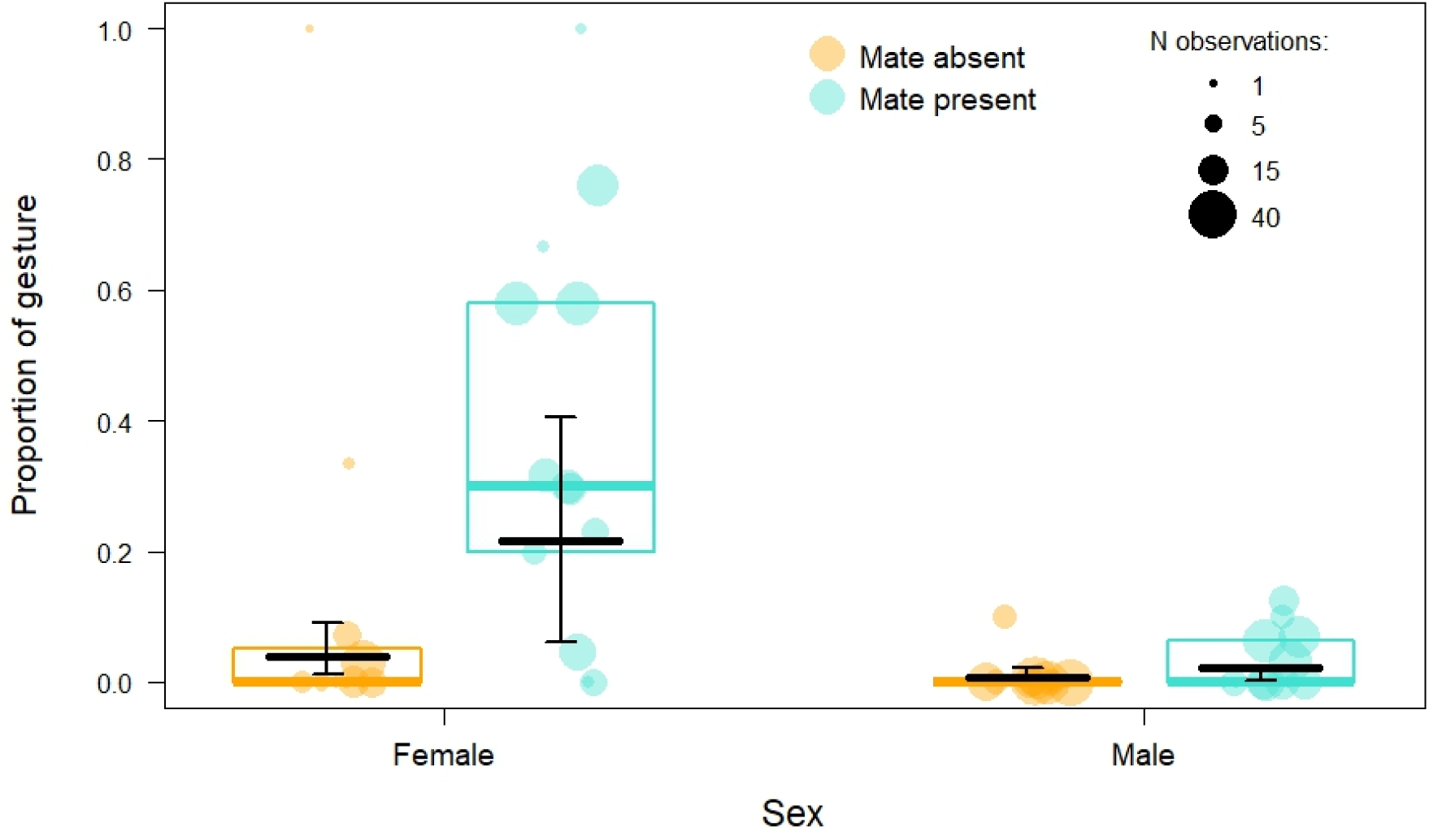
Effect of mate presence on the likelihood of producing a gesture around the nest in male and female great tits. Each dot represents an individual (N = 14 females and 15 males), and the area of the dot is proportional to the number of observations for each individual in each social condition (mate absent in orange and mate present in blue). The boxplot indicates the median and 25% and 75% quartiles across all individuals. The black horizontal line depicts the model prediction derived from Model 1, and the vertical black lines indicate the 89% CI.

**Table 1.**
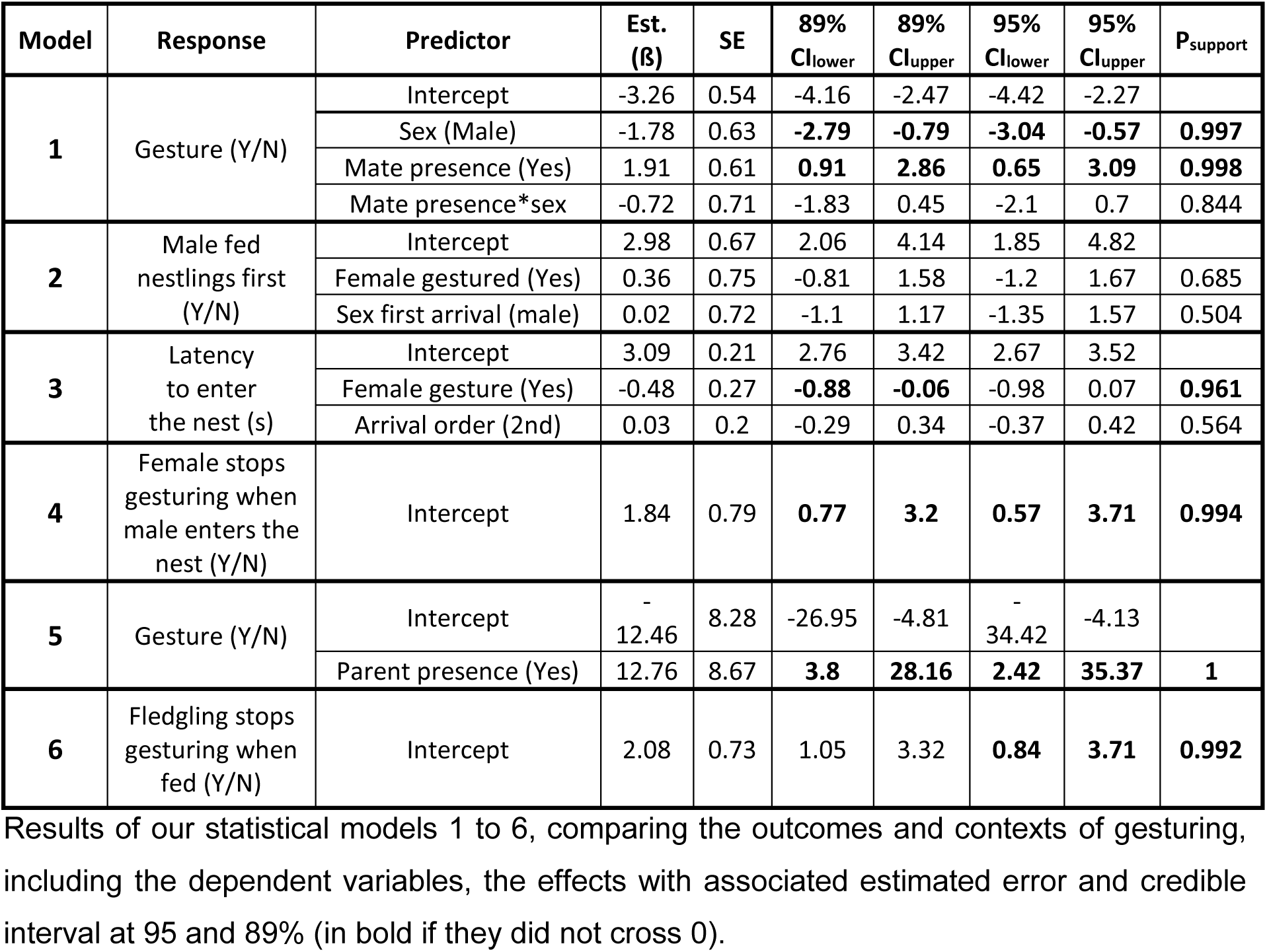
Summary result tables of all the statistical models.

| Model | Response | Predictor | Est. (ß) | SE | 89% Cl<sub>lower</sub> | 89% Cl<sub>upper</sub> | 95% Cl<sub>lower</sub> | 95% Cl<sub>upper</sub> | P<sub>support</sub> |
| --- | --- | --- | --- | --- | --- | --- | --- | --- | --- |
| **1** | Gesture (Y/N) | Intercept | -3.26 | 0.54 | -4.16 | -2.47 | -4.42 | -2.27 |  |
|  |  | Sex (Male) | -1.78 | 0.63 | **-2.79** | **-0.79** | **-3.04** | **-0.57** | **0.997** |
|  |  | Mate presence (Yes) | 1.91 | 0.61 | **0.91** | **2.86** | **0.65** | **3.09** | **0.998** |
|  |  | Mate presence*sex | -0.72 | 0.71 | -1.83 | 0.45 | -2.1 | 0.7 | 0.844 |
| **2** | Male fed nestlings first (Y/N) | Intercept | 2.98 | 0.67 | 2.06 | 4.14 | 1.85 | 4.82 |  |
|  |  | Female gestured (Yes) | 0.36 | 0.75 | -0.81 | 1.58 | -1.2 | 1.67 | 0.685 |
|  |  | Sex first arrival (male) | 0.02 | 0.72 | -1.1 | 1.17 | -1.35 | 1.57 | 0.504 |
| **3** | Latency to enter the nest (s) | Intercept | 3.09 | 0.21 | 2.76 | 3.42 | 2.67 | 3.52 |  |
|  |  | Female gesture (Yes) | -0.48 | 0.27 | **-0.88** | **-0.06** | -0.98 | 0.07 | **0.961** |
|  |  | Arrival order (2nd) | 0.03 | 0.2 | -0.29 | 0.34 | -0.37 | 0.42 | 0.564 |
| **4** | Female stops gesturing when male enters the nest (Y/N) | Intercept | 1.84 | 0.79 | **0.77** | **3.2** | **0.57** | **3.71** | **0.994** |
| **5** | Gesture (Y/N) | Intercept | -12.46 | 8.28 | -26.95 | -4.81 | -34.42 | -4.13 |  |
|  |  | Parent presence (Yes) | 12.76 | 8.67 | **3.8** | **28.16** | **2.42** | **35.37** | **1** |
| **6** | Fledgling stops gesturing when fed (Y/N) | Intercept | 2.08 | 0.73 | 1.05 | 3.32 | **0.84** | **3.71** | **0.992** |

The five adults observed near their nest in natural settings (i.e., not at nestboxes) exhibited wing-fluttering within a 2-meter radius of their nest at least once when their mate was present, after which they entered the natural cavity. One male also displayed wing-fluttering once near the natural nest when alone. These observations are consistent with those made at nestboxes, although males performed the gesture more frequently.

Results of our statistical models 1 to 6, comparing the outcomes and contexts of gesturing, including the dependent variables, the effects with associated estimated error and credible interval at 95 and 89% (in bold if they did not cross 0).

### Social triggers of gesture in adults away from the nest cavity

We found that great tits did not exhibit wing-fluttering when away from the nest cavity (40 females and 38 males). Specifically, males showed no wing-fluttering in 32 observations when alone and in 23 observations when with conspecifics. Females showed no wing-fluttering in 33 observations when alone, and only one observation of wing-fluttering was recorded in 25 observations when with conspecifics (i.e. 4%). In this anecdotal case, the female displayed wing-fluttering when a male approached and subsequently fed her; the behaviour ceased once feeding ended. This observation occurred in the absence of other individuals during a focal period (1 min 55 s). Adults did not perform the gesture when individuals of other species were nearby (N = 5 observations for 5 females and N = 9 observations for 9 males).

### Effect of female gesture on the likelihood for males to enter the nestbox first

In Model 2 (N = 167 observations when both males and females were together at the nest from 22 birds (11 males and 11 females)), we did not find statistical support for an effect of female gesture on the likelihood for males to enter the nestbox first (89% CI largely overlapped 0 and P.support = 0.685). We also found no statistical effect of the order of arrival at the nest on the likelihood for the male to enter the nestbox first (89% CI largely overlapped 0 and P.support = 0.504, Table 1).

### Effect of female gesture on the latency for males to enter the nestbox

In model 3 (N = 112 observations on 11 males), we found a consistent effect of female gesture on the latency for males to enter the nestbox (ß= -0.48, 89% CI [-0.88, -0.06], P.support = 0.961), but with some degree of uncertainty since the 95% CI [-0.98, 0.07] slightly overlapped 0 (Table 1). Males were 1.63 times faster at entering the nestbox when females gestured than when they did not (mean ± SE = 23.77 ± 3.22 sec. without gesture vs. mean ± SE = 14.56 ± 3.17 sec. with gesture, Fig. 2).

**Fig. 2.**
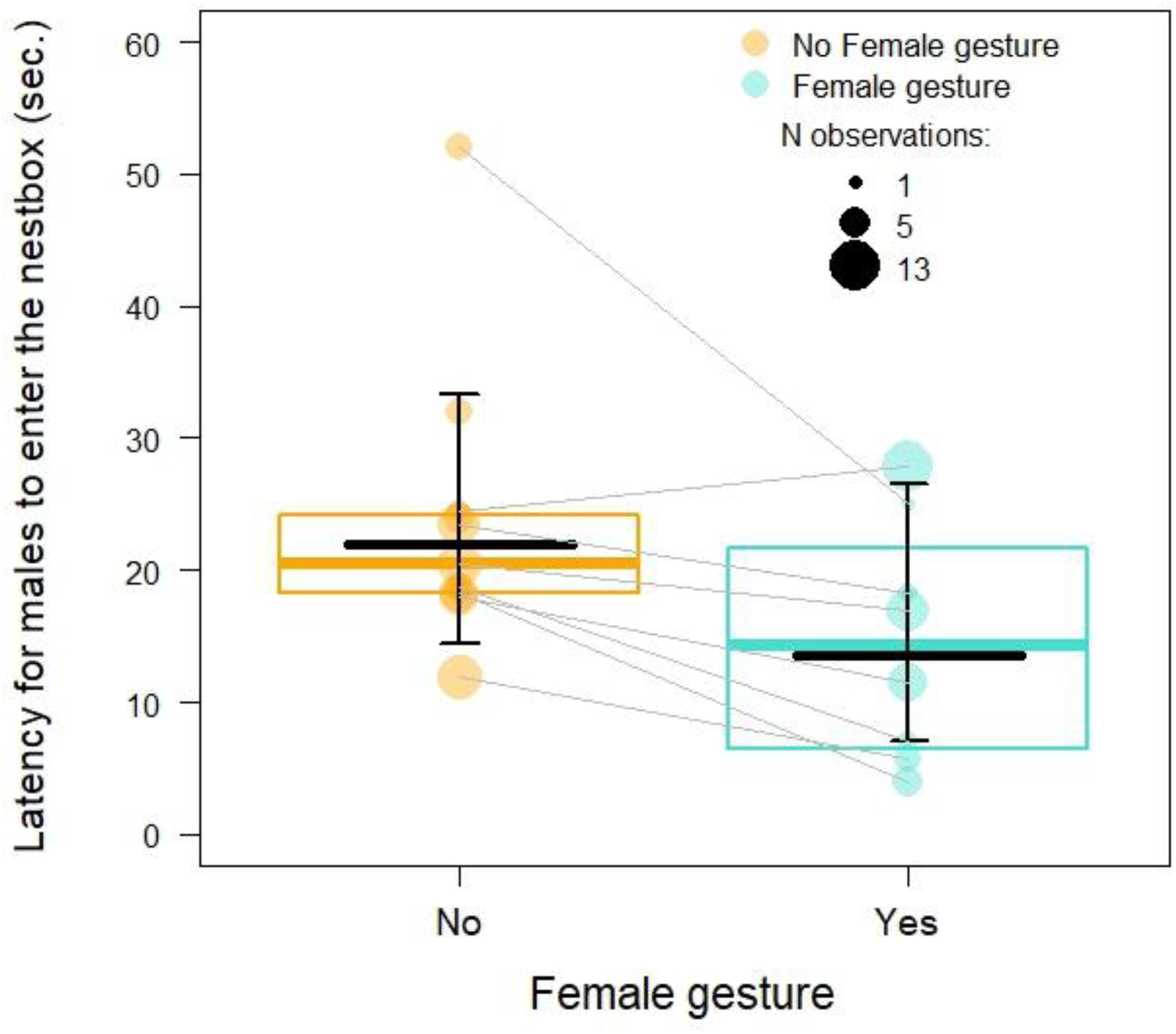
Effect of female gesture on the latency for males to enter the nestbox. Each dot represents an individual (N = 112 observations on 11 males), and the area of the dot is proportional to the number of observations for each male when females did not gesture (orange) and when they produced a gesture (blue). The boxplots indicate the median and 25% and 75% quartiles across all individuals. The black horizontal line depicts the model prediction derived from Model 1a, and the vertical black lines indicate the 89% CI. The thin grey lines link the mean latency for each individual in the absence and presence of females’ gestures (only for the 8 individuals sampled in both conditions).

### Influence of male behaviour on gesture interruption in females at the nestbox

Females stopped gesturing 82.8% of the time when the male entered the nestbox (24 times out of 29 observations of gesture for which we had this information). Said otherwise, females had 4.8 times more chance to stop gesturing than to continue after the male entered the nestbox. Model 4 provides strong statistical support for this pattern (Intercept for female stopping to gesture after the male entered the nestbox: 1.84, 95% CI [0.57, 3.71], *P.support* = 0.994, Table 1). We anecdotally observed two occasions in which a female began wing fluttering while the male had already entered the nest.

### Social influence on fledgling gesture production

The fledglings always produced the wing-fluttering gesture when at least one of the parents was present (36 out of 36 documented gestures). Fledglings gestured in 72% of the observations when parents were present (36 times out of 50 observations) and were never observed gesturing in 49 observations when parents were absent. Our statistical model confirmed the raw observations (effect of parent presence on the likelihood to gesture: ß= 12.76, 95% CI [2.42, 35.37], P.support = 1, Table 1, Fig. 3).

**Fig. 3.**
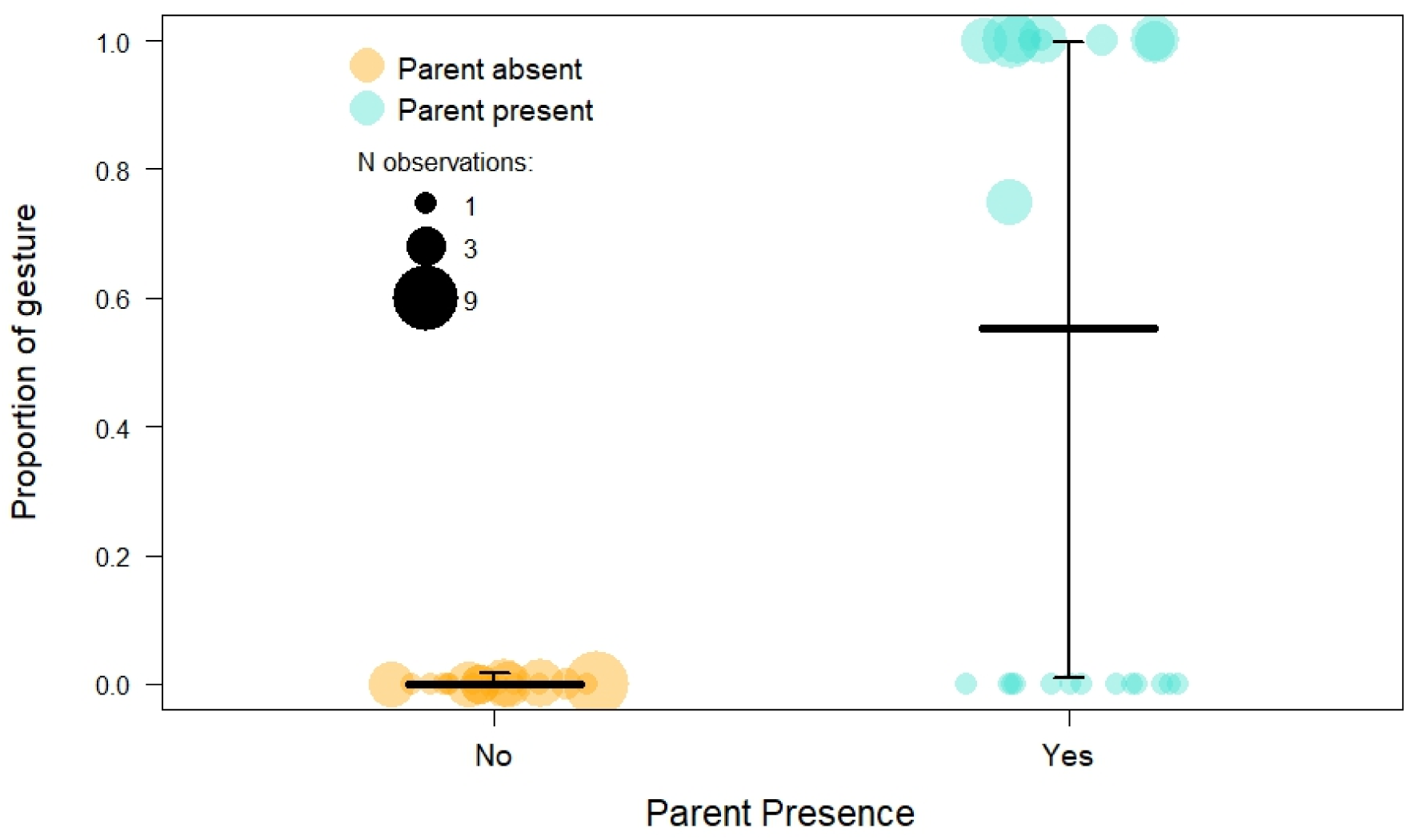
Effect of parent presence on the likelihood of producing a gesture in fledglings. Each dot represents an individual (N = 99 observations on 24 fledglings), and the area of the dot is proportional to the number of observations for each individual in each social condition (parents absent in orange and present in blue). The boxplot indicates the median and 25% and 75% quartiles across all individuals. The black horizontal line depicts the model prediction derived from Model 4, and the vertical black lines indicate the 89% CI.

### Influence of parents’ feeding behaviour on gesture interruption in fledglings

The fledglings stopped gesturing 87.1% of the time when they had been fed by the parent (27 times out of 31 observations of gesture for which we had this information). Said otherwise, fledglings had 6.75 times more chance to stop the gesture than to continue after they had been fed. Our statistical model (Model 6) confirmed the observed pattern (Intercept for fledglings stopping to gesture after being fed: 2.08, 95% CI [0.84, 3.71], *P.support* = 0.999, Table 1).

## Discussion

Research on complex gestures in animals outside primates remains rare despite their crucial importance in understanding the evolution of complex communication, including language. Here, through observations of both juvenile and adult birds in natural ecological settings, we show that the wing-fluttering behaviour of great tits could meet the criteria for symbolic gestures; (1) it is produced primarily in the presence of the gesture’s intended target (mates for adults, and parents for fledglings), (2) it elicits a specific response and (3) stops once the goal has been reached, and (4) does not act as a direct physical agent. These findings mirror previous observations in adult Japanese tits (Suzuki and Sugita, 2024). In addition, our study extends this work by examining the functional use of this gesture in fledglings, revealing a developmental shift in its function between fledglings and adults.

First, great tits exhibit wing-fluttering behaviour only in the presence of the target receiver (i.e. mates for adults and parents for fledglings), and specifically in a feeding context. Fledglings never displayed this gesture when they were only with their siblings from the same brood (i.e. when no parents were present), and adults did not perform this gesture when they were without their mate, even if individuals of other species were nearby. Second, this behaviour elicits a specific response from the receiver. Although the females’ gesture has no effect on the likelihood of males entering the nest box first, males tend to be faster when the gesture is made. Similar to what has been reported in Japanese tits (Suzuki and Sugita, 2024), we observed that females exhibited wing-fluttering more frequently than males. However, in our population, female great tits feed the nestlings less often than males, and this display does not seem to correspond to the “after you” signal described in Japanese tits (Suzuki and Sugita, 2024), but rather could convey a “go ahead” message. Third, we found that the emitter stops gesturing once he has achieved the desired outcome (= the receiver has produced a specific behaviour in response to the signal); adults were significantly more likely to stop wing fluttering after their partner entered the nestbox, and fledglings stopped wing-fluttering once their parents fed them. Finally, the fourth criterion of symbolic gesturing is also fulfilled: the emitter never acts as a direct physical agent. In adults, wing-fluttering elicits nest entry from the mate without physical contact, and in juveniles, it prompts food delivery without direct interaction. Taken together, these results indicate some evidence of intentional communication (Townsend et al., 2017; Aychet et al., 2025; Graham, 2025). It will also be important to rule out arousal-based explanations for this wing-fluttering behaviour (Graham, 2025). For instance, in the case of re-caching and “Theory of mind” in jays, the stress induced by the presence of another individual can increase arousal levels and alter caching behaviour (Van Der Vaart et al., 2012). Thus, what appears to be a cognitively sophisticated response may, in fact, be explained by simpler stress-related mechanisms. In our case, if waiting outside the nest or if the presence of the mate near the nest is stressful, it is possible that wing-fluttering represents a stress response that delays or inhibits the individual’s entry. Hormonal studies could help assess the likelihood of arousal-based explanations for these signals. However, the response of the mate to wing fluttering is unlikely to be related to stress, and a purely stress-based account does not readily explain why males enter the nest more quickly following female wing-fluttering than when the gesture is absent. Females do not systematically perform wing-fluttering whenever the mate is present, suggesting that the behaviour is not merely a by-product of social stress. Together, these observations indicate that while stress-related arousal may contribute to the expression of wing-fluttering, it is unlikely to fully account for its functional role, leaving open the possibility that the gesture serves a communicative function.

Our results reveal how fledglings use wing-fluttering to signal to their parents that they want to be fed, and how, later in life, adult tits employ the same gesture to prompt their mates to enter the nest cavity and feed the nestlings, highlighting its remarkable flexibility in both acquisition and use. Our results suggest that adult tits employ wing-fluttering in a triadic interaction framework, in which a female signal (the emitter) prompts the receiver (the male) to act on a third party (the nestlings), aiming at a specific goal (nest entry). In contrast, fledglings exhibit this behaviour solely in dyadic interactions, where the emitter of the signal directly benefits from the response of the receiver. The triadic interaction observed here contrasts with most gestures in apes, which are typically dyadic and involve engaging in dyadic interaction (e.g., approaching, grooming, travelling together, climbing on the back, playing; Tomasello and Call, 2019), whereas in tits the male does not behaviourally act back on the female, but they coordinate their actions with respect to a third party. Fledglings may innately perform this gesture as an impulsive response to the excitement of a parent’s arrival, aiming to attract attention and be fed. At this stage, the behaviour may be uncontrolled and unintentional. However, it is possible that, over time, individuals learn to regulate and refine this movement, eventually using it purposefully to achieve specific communicative goals; they will employ wing-fluttering not only as a symbolic gesture but also within a triadic context (i.e. one involving a signaller, a receiver, and a specific goal) highlighting a more complex form of gestural communication. The intentional use of gestures may therefore develop over the course of ontogeny, as observed in chimpanzee and human infants, where markers of intentional communication increase with age (chimpanzees: Bard et al., 2014; Fröhlich et al., 2019; humans: Bates et al., 1979; Carpenter et al., 1998). Longitudinal studies tracking the development of gesture and intentionality from birth may offer valuable insights into how intentional complex gestural communication in a triadic context emerges, not only in primates (Liebal et al., 2019), but also in other species such as birds.

We found that wing-fluttering behaviour was exhibited in a feeding/foraging context. It would be interesting to investigate whether this gesture is also used in other social contexts. Playback experiments reported that birds can display wing-fluttering during mobbing events (Salis et al., 2023; Coye and Dutour, 2025), when individuals collectively mob a predator to drive it away. Given the inherently social nature of mobbing - involving multiple individuals - it raises the possibility that wing-fluttering may serve as a signal from one prey individual to others, potentially to prompt action or coordination (e.g., “you go first”). Such use would again imply a triadic interaction framework, involving an emitter, a receiver, and a shared goal (i.e., the predator). Moreover, great tits are known to combine calls into their mobbing sequences using syntactic rules (Dutour et al., 2019a). This makes the mobbing context particularly relevant for investigating the co-evolution of visual and vocal communication systems. Studying how these modalities interact could provide new insights into the evolutionary precursors of language. Future experiments combining playbacks and predator presentations would be necessary to explore these hypotheses further.

Research over the past ten years has discovered some rudimentary but fundamental building blocks of human language in Japanese and great tits. Both species of tits combine meaningful calls to form meaningful call combinations that can be seen as rudimentary compositional syntax (Suzuki et al., 2016; Dutour et al., 2019a). Our current study, together with the recent findings from Suzuki and Sugita (2024) on Japanese tits, suggest the presence of symbolic gestures in tits, which are also argued to be a fundamental building block that could have led to the evolution of language. Combinatorial and symbolic communication are two fundamental aspects of human language, placing the tits as an enlightening study species for comparative studies aimed at retracing the evolutionary origins of core fundamental principles of language (Salis et al., 2025). The current knowledge, reporting a single gesture and a single call combination in tits indicates that these capacities might be much more rudimentary or limited than in great apes. Chimpanzee for instance express a large range of messages across diverse social situation using a communicative system that integrate both highly versatile vocal combinations (Bortolato et al., 2023; Girard-Buttoz et al., 2022; Girard-Buttoz et al., 2025) and a large repertoire of gestures (Hobaiter and Byrne, 2014). Future studies should embrace the challenge of recording gestural and vocal communication of tits across all daily life situations to establish if this striking difference with apes is real or only reflects methodological constraints.

Our findings suggest that great tits use wing-fluttering as a gesture in dyadic contexts during the first months of life, and later, as adults, use this gesture in triadic contexts involving a signaller (the female), a receiver (the male), and a specific goal (nest entry). This developmental shift highlights the flexibility of gestural communication and supports the idea that such complex gestures are not limited to primates. Unlike most previous studies conducted in controlled or captive environments, our study was carried out in natural settings, allowing us to account for ecological pressures that shape communication. Finally, future research on both visual and acoustic communication in both Paridae and non-Paridae species and across other animal groups will be essential to deepen our understanding of the evolutionary roots of complex communication systems, including those underlying human language.

### Ethics

All protocols were in accordance with current French laws and adhered to the Guidelines for the Use of Animals of the Association for the Study of Animal Behaviour/Animal Behavior Society. We conducted only observational work on great tits in their natural environment, without any experimental manipulation.

## Data availability

The data used for this study are available in the GitHub repository (https://github.com/tozbu/tit_gesture).

## Code availability

The R codes used for this study are available in the repository (https://github.com/tozbu/tit_gesture).

## Acknowledgements

We thank Rose Beghin, Suzanne Busquet-Losserand, Candice Heluwaert, Maé Leclézio, and Killian Thollot for fieldwork assistance. We thank the Pierre Vérots Foundation for granting access and supporting the establishment of the population ten years ago.

## Author contributions

M.D.: conceptualization, supervision, visualization, data collection, data curation, formal analysis, investigation, methodology, project administration, visualization, writing – original draft; V.O.: methodology, data collection, writing – review and editing; T.L.: conceptualization, methodology, funding acquisition, writing – review and editing; C.G.-B.: conceptualization, formal analysis, writing – original draft.

## References

1. Aychet, J., Dafreville, M., & Canteloup, C. (2025). Intentional Communication in Animals.

2. Bard, K. A., Dunbar, S., Maguire-Herring, V., Veira, Y., Hayes, K. G., & McDonald, K. (2014). Gestures and social-emotional communicative development in chimpanzee infants. American Journal of Primatology, 76(1), 14–29.

3. Bortolato, T., Friederici, A. D., Girard-Buttoz, C., Wittig, R. M., & Crockford, C. (2023). Chimpanzees show the capacity to communicate about concomitant daily life events. Iscience, 26(11).

4. Bates, E., Benigni, L., Bretherton, I., Camaioni, L., & Volterra, V. (1979). The emergence of symbols: Cognition and communication in infancy. New York: Academic Press.

5. Buerkner, P. C. (2018). Advanced Bayesian Multilevel Modeling with the R Package brms. The R Journal, 10(1), 395–411. doi:10.32614/RJ-2018-017

6. Carpenter, M., Nagell, K., Tomasello, M., Butterworth, G., & Moore, C. (1998). Social cognition, joint attention, and communicative competence from 9 to 15 months of age. Monographs of the society for research in child development, i-174.

7. Coye, C., & Dutour, M. (2025). Mobbing sequences of American wrens elicit mobbing responses in European tits. Animal Behaviour, 221, 123050.

8. Da Silva, S., & Matsushita, R. (2025). Feathers of Grace: The “After You” Gesture in Japanese Tits. Biology, 14(3), 297.

9. Dutour, M., Léna, J. P., & Lengagne, T. (2017). Mobbing calls: a signal transcending species boundaries. Animal Behaviour, 131, 3–11.

10. Dutour, M., Lengagne, T., & Léna, J. P. (2019a). Syntax manipulation changes perception of mobbing call sequences across passerine species. Ethology, 125(9), 635–644.

11. Dutour, M., Léna, J. P., Dumet, A., Gardette, V., Mondy, N., & Lengagne, T. (2019b). The role of associative learning process on the response of fledgling great tits (*Parus major*) to mobbing calls. Animal Cognition, 22, 1095–1103.

12. Eleuteri, V., Bates, L., Nyaradzo Masarira, Y., Plotnik, J. M., Hobaiter, C., & Stoeger, A. S. (2025). Investigating intentionality in elephant gestural communication. Royal Society Open Science, 12(7).

13. Fröhlich, M., Wittig, R. M., & Pika, S. (2019). The ontogeny of intentional communication in chimpanzees in the wild. Developmental science, 22(1), e12716.

14. Girard-Buttoz, C., Zaccarella, E., Bortolato, T., Friederici, A. D., Wittig, R. M., & Crockford, C. (2022). Chimpanzees produce diverse vocal sequences with ordered and recombinatorial properties. Communications Biology, 5(1), 410.

15. Girard-Buttoz, C., Neumann, C., Bortolato, T., Zaccarella, E., Friederici, A. D., Wittig, R. M., & Crockford, C. (2025). Versatile use of chimpanzee call combinations promotes meaning expansion. Science Advances, 11(19), eadq2879.

16. Goodwyn, S. W., Acredolo, L. P., & Brown, C. A. (2000). Impact of symbolic gesturing on early language development. Journal of Nonverbal behavior, 24, 81–103.

17. Graham, K. E. (2025). Goal-directed bodily signals in birds and frogs. Learning & Behavior, 53(1), 5–6.

18. Harding, C. (1984). Acting with intention: A framework for examining the development of the intention to communicate. The origins and growth of communication, 123–135.

19. Hobaiter, C., & Byrne, R. W. (2014). The meanings of chimpanzee gestures. Current Biology, 24(14), 1596–1600.

20. Kalan, A. K., Nakano, R., & Warshawski, L. (2025). What we know and don’t know about great ape cultural communication in the wild. American Journal of Primatology, 87(1), e23560.

21. Leavens, D. A., Russell, J. L., & Hopkins, W. D. (2005). Intentionality as measured in the persistence and elaboration of communication by chimpanzees (*Pan troglodytes*). Child development, 76(1), 291–306.

22. Liebal, K., Schneider, C., & Errson-Lembeck, M. (2019). How primates acquire their gestures: evaluating current theories and evidence. Animal Cognition, 22(4), 473–486.

23. Liszkowski, U., Carpenter, M., Henning, A., Striano, T., & Tomasello, M. (2004). Twelve-month-olds point to share attention and interest. Developmental science, 7(3), 297–307.

24. Moore, R. (2016). Meaning and ostension in great ape gestural communication. Animal Cognition, 19(1), 223–231.

25. Moura, L. N., Silva, M. L., Garotti, M. M., Rodrigues, A. L., Santos, A. C., & Ribeiro, I. F. (2014). Gestural communication in a new world parrot. Behavioural processes, 105, 46–48.

26. Pika, S., & Bugnyar, T. (2011). The use of referential gestures in ravens (*Corvus corax*) in the wild. Nature communications, 2(1), 560.

27. Pika, S., Liebal, K., Call, J., & Tomasello, M. (2005). Gestural communication of apes. Gesture, 5(1-2), 41–56.

28. Pollick, A. S., & de Waal, F. B. (2007). Ape gestures and language evolution. Proceedings of the National Academy of Sciences, 104(19), 8184–8189.

29. Salis, A., Lena, J.-P., & Lengagne, T. (2022). Which acoustic parameters modify the great tit’s response to conspecific combinatorial mobbing calls? Behavioral Ecology and Sociobiology, 76(4), 46. 10.1007/s00265-022-03157-x

30. Salis, A., J.-P. Léna, and T. Lengagne. (2023). Both Learning and Syntax Recognition Are Used by Great Tits When Answering to Mobbing Calls. Behavioral Ecology, 34(6), arad061. 10.1093/beheco/arad06

31. Salis, A., Ryder, R. J., Molina, A., Schlenker, P., & Chemla, E. (2025). Birds combined calls more than 11 million years ago. Scientific Reports, 15(1), 21338.

32. Schlenker, P., Salis, A., Leroux, M., Coye, C., Rizzi, L., Steinert-Threlkeld, S., & Chemla, E. (2024). Minimal Compositionality versus Bird Implicatures: two theories of ABC-D sequences in Japanese tits. Biological Reviews, 99(4), 1278–1297.

33. Stan Development Team (2020). RStan: the R interface to Stan. R package version 2.21.2. https://mc-stan.org/

34. Suzuki, T. N., & Sugita, N. (2024). The ‘after you’ gesture in a bird. Current Biology, 34(6), R231–R232.

35. Suzuki, T. N., & Matsumoto, Y. K. (2022). Experimental evidence for core-Merge in the vocal communication system of a wild passerine. Nature Communications, 13(1), 5605.

36. Suzuki, T. N., Wheatcroft, D., & Griesser, M. (2016). Experimental evidence for compositional syntax in bird calls. Nature communications, 7(1), 10986.

37. Tomasello, M., & Call, J. (2019). Thirty years of great ape gestures. Animal cognition, 22(4), 461–469.

38. Townsend, S. W., Koski, S. E., Byrne, R. W., Slocombe, K. E., Bickel, B., Boeckle, M., … & Manser, M. B. (2017). Exorcising G rice’s ghost: An empirical approach to studying intentional communication in animals. Biological Reviews, 92(3), 1427–1433.

39. Vail, A. L., Manica, A., & Bshary, R. (2013). Referential gestures in fish collaborative hunting. Nature communications, 4(1), 1765.

40. Van Der Vaart, E., Verbrugge, R., & Hemelrijk, C. K. (2012). Corvid re-caching without ‘theory of mind’: A model. PloS one, 7(3), e32904.

