## Supplementary Material for "Ontogenic shift in the usage and function of a symbolic bird gesture"

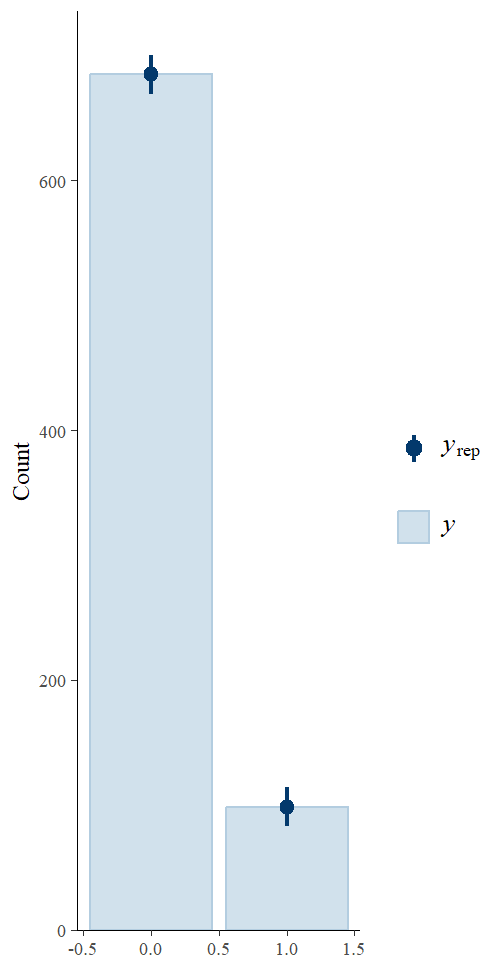

**Fig. S1. Posterior predictive check for Model 1.**

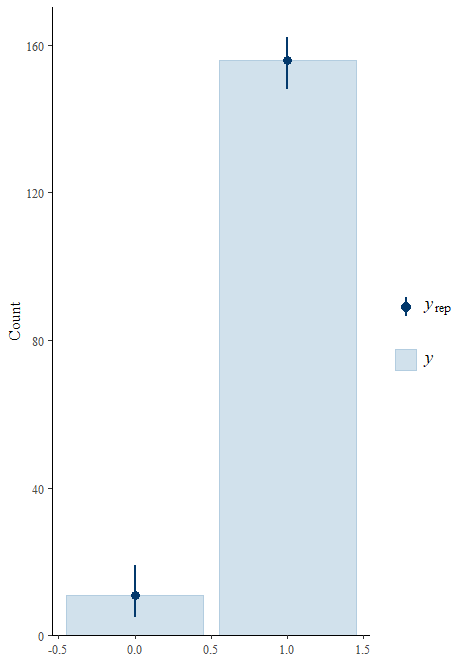

**Fig. S2. Posterior predictive check for Model 2.**

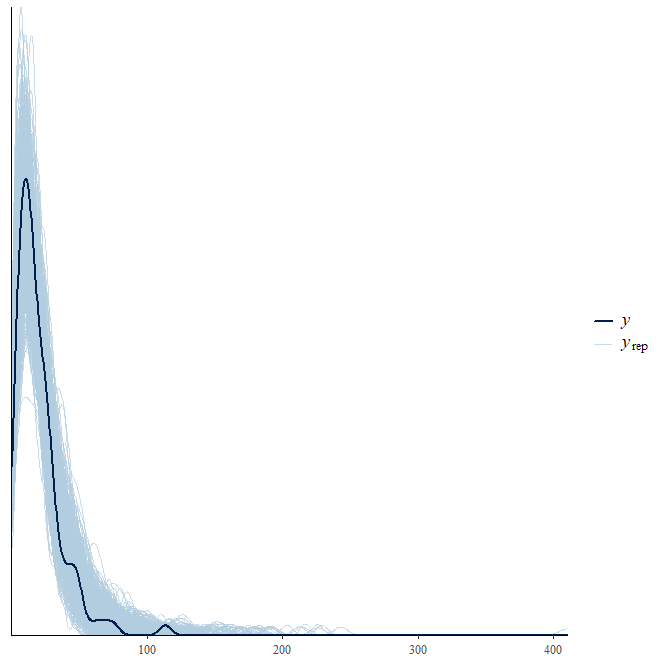

**Fig. S3. Posterior predictive check for Model 3.**

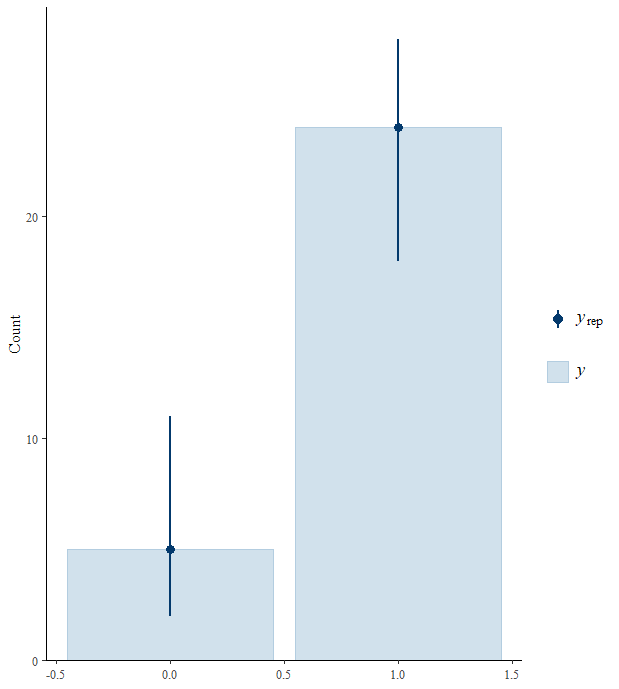

**Fig. S4. Posterior predictive check for Model 4.**

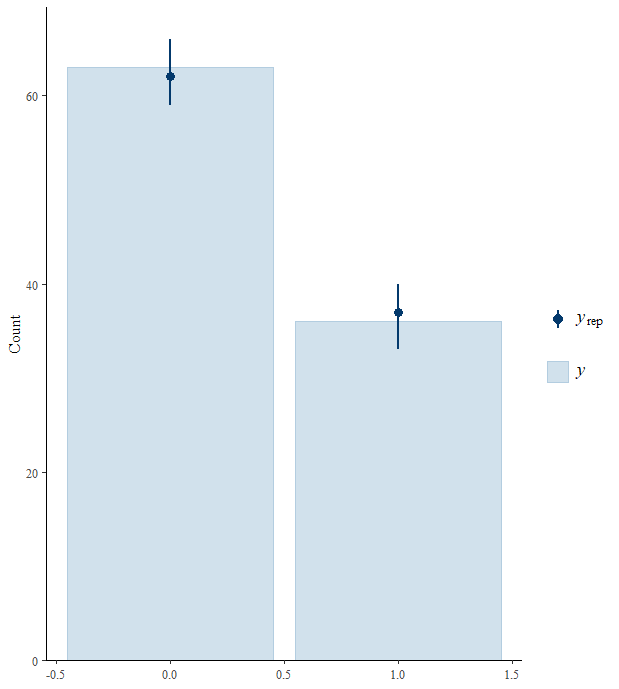

**Fig. S5. Posterior predictive check for Model 5.**

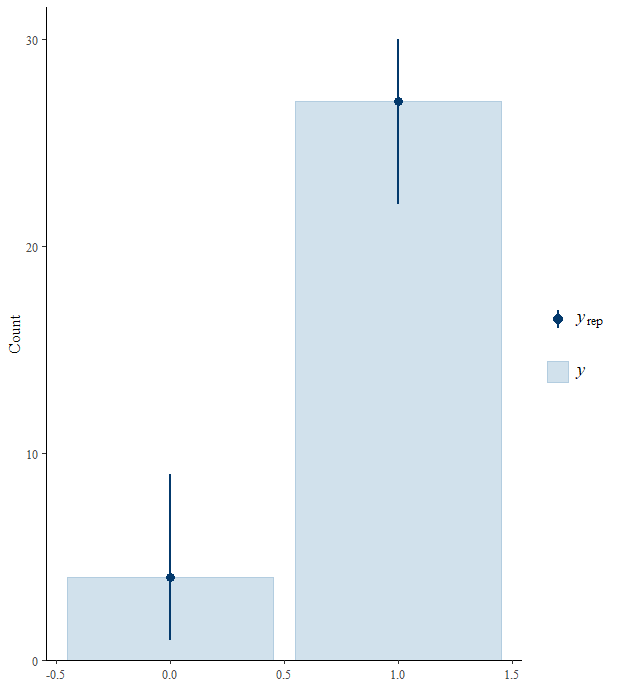

**Fig. S6. Posterior predictive check for Model 6.**
